# A Novel Initiation Pathway in *Escherichia Coli* Transcription

**DOI:** 10.1101/042432

**Authors:** Eitan Lerner, SangYoon Chung, Benjamin L. Allen, Shuang Wang, Jookyung J. Lee, Winson S. Lu, Wilson L. Grimaud, Antonino Ingargiola, Xavier Michalet, Yazan Alhadid, Sergei Borukhov, Terence Strick, Dylan J. Taatjes, Shimon Weiss

**Affiliations:** Dept. of Chemistry & Biochemistry, University of California Los Angeles, Los Angeles, CA 90095.; Dept. of Chemistry & Biochemistry, University of Colorado, Boulder.; Institut Jacques Monod, Centre National de la Recherche Scientifique and University of Paris Diderot and Sorbonne Paris Cité, Paris, France.; Rowan University School of Osteopathic Medicine, Stratford, NJ 08084, USA.; California NanoSystems Institute, University of California Los Angeles, Los Angeles, CA 90095.; Dept. of Physiology, University of California Los Angeles, Los Angeles, CA 90095.

**Keywords:** DNA transcription, transcription initiation, pausing, backtracking, kinetics, single molecule, FRET, magnetic tweezers

## Abstract

Initiation is a highly regulated, rate-limiting step in transcription. We employed a series of approaches to examine the kinetics of RNA polymerase (RNAP) transcription initiation in greater detail. Quenched kinetics assays, in combination with magnetic tweezer experiments and other methods, showed that, contrary to expectations, RNAP exit kinetics from later stages of initiation (e.g. from a 7-base transcript) was markedly slower than from earlier stages. Further examination implicated a previously unidentified intermediate in which RNAP adopted a long-lived backtracked state during initiation. In agreement, the RNAP-GreA endonuclease accelerated transcription kinetics from otherwise delayed initiation states and prevented RNAP backtracking. Our results indicate a previously uncharacterized RNAP initiation state that could be exploited for therapeutic purposes and may reflect a conserved intermediate among paused, initiating eukaryotic enzymes.

**Significance:** Transcription initiation by RNAP is rate limiting owing to many factors, including a newly discovered slow initiation pathway characterized by RNA backtracking and pausing. This backtracked and paused state occurs when all NTPs are present in equal amounts, but becomes more prevalent with NTP shortage, which mimics cellular stress conditions. Pausing and backtracking in initiation may play an important role in transcriptional regulation, and similar backtracked states may contribute to pausing among eukaryotic RNA polymerase II enzymes.

## Introduction

Transcription of genomic DNA requires formation of an RNA polymerase (RNAP)-promoter initially-transcribing complex (RP_ITC_), in which RNAP unwinds DNA and generates a mechanically-stressed intermediate (1) through DNA scrunching (2, 3). In transcription initiation, the interaction of the σ70 subunit of RNAP with the promoter creates a physical barrier for the nascent RNA transcript (4, 5). For productive transcription to take place this barrier must be removed (4, 6), otherwise RNAP will enter repeated cycles of unsuccessful transcription attempts known as abortive initiation (7, 8).

When RNAP experiences multiple cycles of abortive initiation, release of the nascent RNA is rate limiting for each cycle (3, 9). The release of an abortive transcript is achieved by consecutive backward DNA translocations, in which the short RNA transcript backtracksthrough the secondary channel until it is released.

To study the mechanism of transcription initiation in greater detail, we developed a solution-based, single-run quenched kinetics transcription assay that measures the kinetics of run-off RNA products. Using this assay we assessed the kinetics out of NTP-starved RP_ITC_ states using an E. coli transcription system reconstituted out of wt RNAP subunits. We also performed quenched kinetics assays using standard in vitro approaches to examine abortive initiation in various contexts, including NTP starvation. Lastly, we performed single-molecule magnetic tweezers experiments to monitor temporal trajectories of RNAP-DNA complexes during transcription initiation.

## Results and Discussion

### Single-round transcription quenched kinetics assay

To quantitatively study the mechanism of transcription initiation by RNAP, we developed a single-round quenched kinetics assay (**Fig. 1**) to probe the kinetics of transcription by directly counting the number of transcripts produced over time. Using this assay, we examined the kinetics of *E. coli* RNAP transcription from distinct RP_ITC_ states generated via NTP starvation (**Figs. 1A** & **1B**). Because transcription initiation is much slower than elongation (i.e. initiation is rate-limiting) (9), the synthesis of relatively short, yet fulllength RNA products (39-and 41-base transcripts) reflects the rate of transcription initiation (**Figs. 1C** & **1D**).

**Fig. 1:**
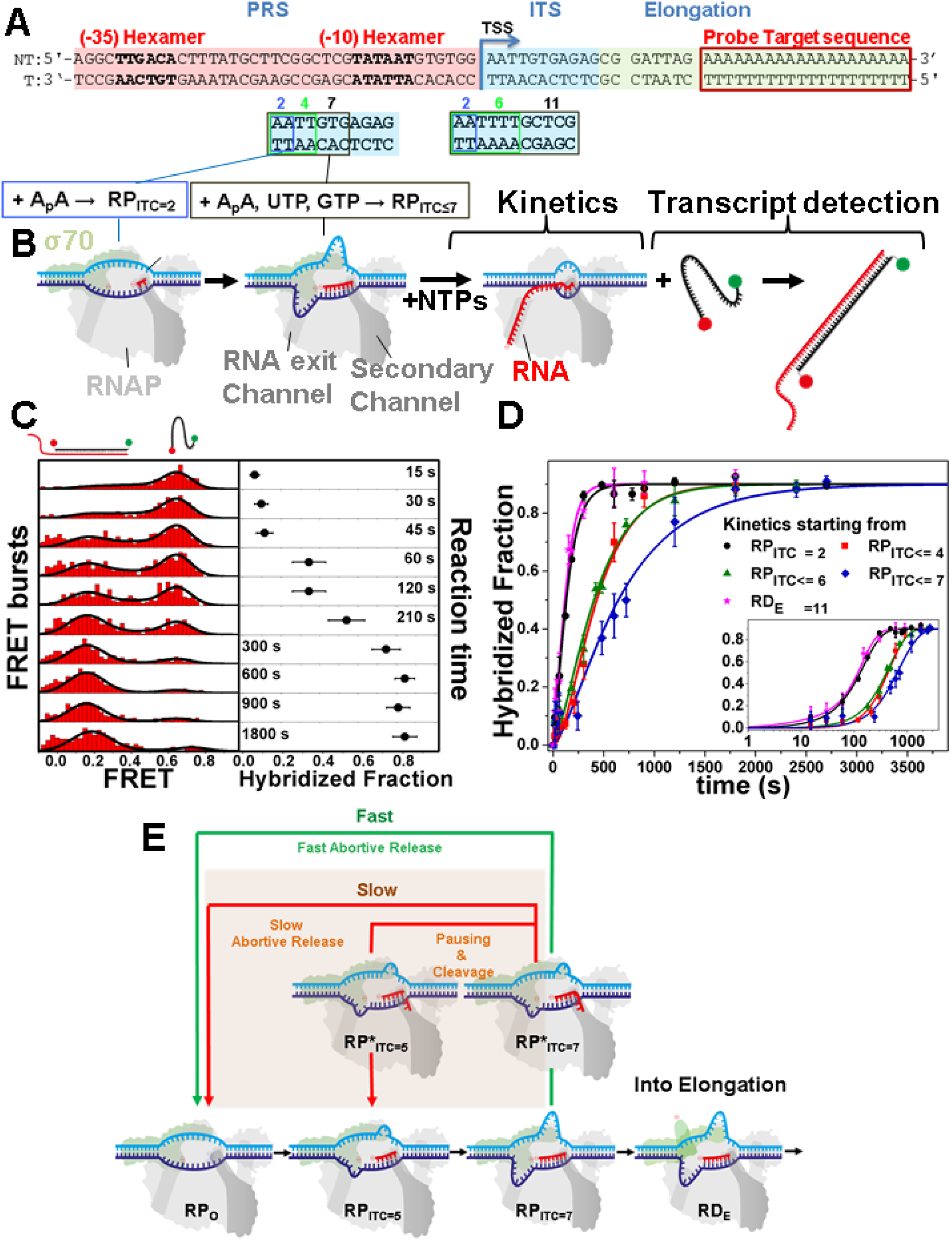
Quenched kinetics transcription results identify an initiation-related stalled state. Representative promoter sequence used here to show how by changing the initially transcribed sequence (ITS, cyan), different NTP-starved states can be generated (RP_ITC=2_, RP_ITC≤4, 6, 7_, RD_E=11_). Other regions of the promoter include the promoter recognition sequence (PRS, pink) and the elongation sequence (yellow), including a probe target complementary sequence (red). All promoters measured are described in **Fig. S1**. (B) Schematic of RNAP run-off transcription starting from a particular NTP-starved state. Upon supplementing all NTPs, transcription kinetics starts and transcripts are quantified via hybridization to a FRET probe. (C) Example of quenched kinetics data generated from quantification of run-off transcripts. (D) Run-off kinetics from various NTP-starved states. Kinetics starting from late initiation states (e.g. RP_ITC≤7_, blue) are slower than from an earlier initiation state (e.g. RP_ITC=2_, black). (E) A schematic of RNAP transcription initiation based upon classical models and the novel intermediate described here. We identify a slower initiation pathway (highlighted red) involving RNAP backtracking, in which backtracked RNA is ultimately cleaved or abortively released.

The quenched kinetics transcription assay is based on quantification of single run-off transcripts by hybridization with a doubly-labeled ssDNA probe (**Fig. 1B**). The number of hybridized probes (and hence the number of transcripts) is accurately determined using μs alternating-laser excitation (μsALEX)-based fluorescence-aided molecule sorting (ALEX-FAMS) (10, 11) (**Fig. 1C**). ALEX-FAMS is a method based on single-molecule Förster Resonance Energy Transfer (smFRET) (12). A significant advantage of smFRET and ALEX-FAMS is their ability to identify distinct populations in a model-free manner, simply by counting single-molecule events of one sort (with a given FRET efficiency population) and comparing their number to the count of single-molecule events of another sort (representing a distinct FRET efficiency population) (10, 11, 13, 14) (**Fig. 1C**). Hence, FAMS is a suitable method for the quantification of run-off transcripts at picomolar probe concentrations. At such low RNAP-promoter concentrations, the time for re-association of σ70 with the RNAP core enzyme and the time for re-association of the formed holoenzyme with the promoter are longer than the time for a single transcription run; therefore, single-run reaction conditions are achieved. We designed the transcribed DNA sequence so that the probe hybridization sequence was at the end of the transcript (**Fig. 1A**). Therefore, probe hybridization will not interfere with transcription initiation.

The ssDNA probe was doubly-labeled with a FRET pair. When free in solution, the probe yields a single FRET population with peak FRET efficiency of E~0.75. Hybridization with the run-off RNA transcript yields a FRET efficiency population with a lower peak value of E~0.3, due to the probe being stretched via hybridization to the 20A target sequence segment of the run-off transcript. Using this assay to assess transcription initiation kinetics required: (i) formation of a stable initial state, (ii) addition of NTPs at zero timepoint, (iii) efficiently and rapidly quenching the reaction at selected times, (iv) promotion of full hybridization of the ssDNA FRET probe to transcripts, and (v) prevention of RNA degradation.

We designed the initial transcription sequence (ITS) of the lacCONS (15, 16) (**Fig. 1A**) or T5N25(3) (**Fig. S1**) promoters so that RNAP would transcribe abortive products of varying maximal lengths upon addition of a partial set of NTPs (**Fig. S1**). Run-off transcript production kinetics, starting from an NTP-starved state, were measured using a constant t_entrance_ and varying t_exit_ incubation times. Stable open complexes were formed by adding an initiating dinucleotide (A_p_A or A_p_U in the case of lacCONS or T5N25 promoter, respectively) to achieve RP_ITC=2_. To stabilize an RP_ITC_ state (RP_ITC≤i_, i e [4,6,7]) or a RNAP-DNA Elongation (RD_E=11_) complex, RP_ITC=2_ was incubated for a given time, t_entrance_, in the presence of a partial set of NTPs. The missing NTPs were then added and the system was incubated for another time period, t_exit_ (**Fig. 1B**). The transcription reaction was then quenched by adding 0.5 M guanidine hydrochloride (GndHCl). GndHCl served both as a reaction quencher (**Fig. S2**) and as an enhancer of hybridization of the ssDNA FRET probe to the run-off transcript (**Fig. S3**). After quenching the transcription reaction, the 20-base ssDNA FRET probe was added, followed by FAMS-ALEX measurements to determine the number of run-off transcripts per time point (**Fig. 1C**).

### Slower transcription initiation kinetics from select NTP starved states

We anticipated that RNAP transcription kinetics from a late RP_ITC≤i_ state (‘exit kinetics’) would be similar, if not faster, than from an early RP_ITC≤j_ state (i>j). Strikingly, however, we observed that exit kinetics from the RP_ITC≤4_, RP_ITC≤6_, or RP_ITC≤7_ states were slower than from RP_ITC=2_ state (**Fig. 1D**). In fact, whereas exit kinetics from RP_ITC=2_ was similar to that of RNAP already in the elongation state (RD_E=11_), exit kinetics from RP_ITC≤7_ was at least 3.5-fold slower (**Table S1**). These results suggest that, contrary to existing models, a previously undetected state exists for RNAP initiation complexes (RP*_ITCC≤i_, **Fig. 1E**). Importantly, this state was transient and overall RNAP activity remained unchanged, given that all “delayed” RNAP complexes (RP_ITC≤4_, RP_ITC≤6_, or RP_ITC≤7_) eventually transitioned to elongation (**Fig. 1D**).

### Delayed initiation kinetics associated with backtracking

It is well-established that elongating RNAP enzymes can backtrack and pause (17, 18). In such circumstances, the nascent RNA 3’-end backtracks into the secondary channel, where it is subject to endonucleolytic cleavage by the GreA-RNAP complex (17, 19, 20). To test whether delayed exit kinetics for RP_ITC≤7_ was due to backtracking during initiation, we assessed the effect of GreA using our single-round quenched kinetics assay. As shown in **Figs. 2A & 2B**, 1 μM GreA accelerated the exit kinetics from RP_ITC≤7_ relative to the exit kinetics from RP_ITC=2_. (~50% recovery from the RP*_ITC≤7_ delayed state; see Methods and **Table S1**). These results are consistent with GreA-dependent release of RNAP from a backtracked and paused state during initiation.

**Fig. 2:**
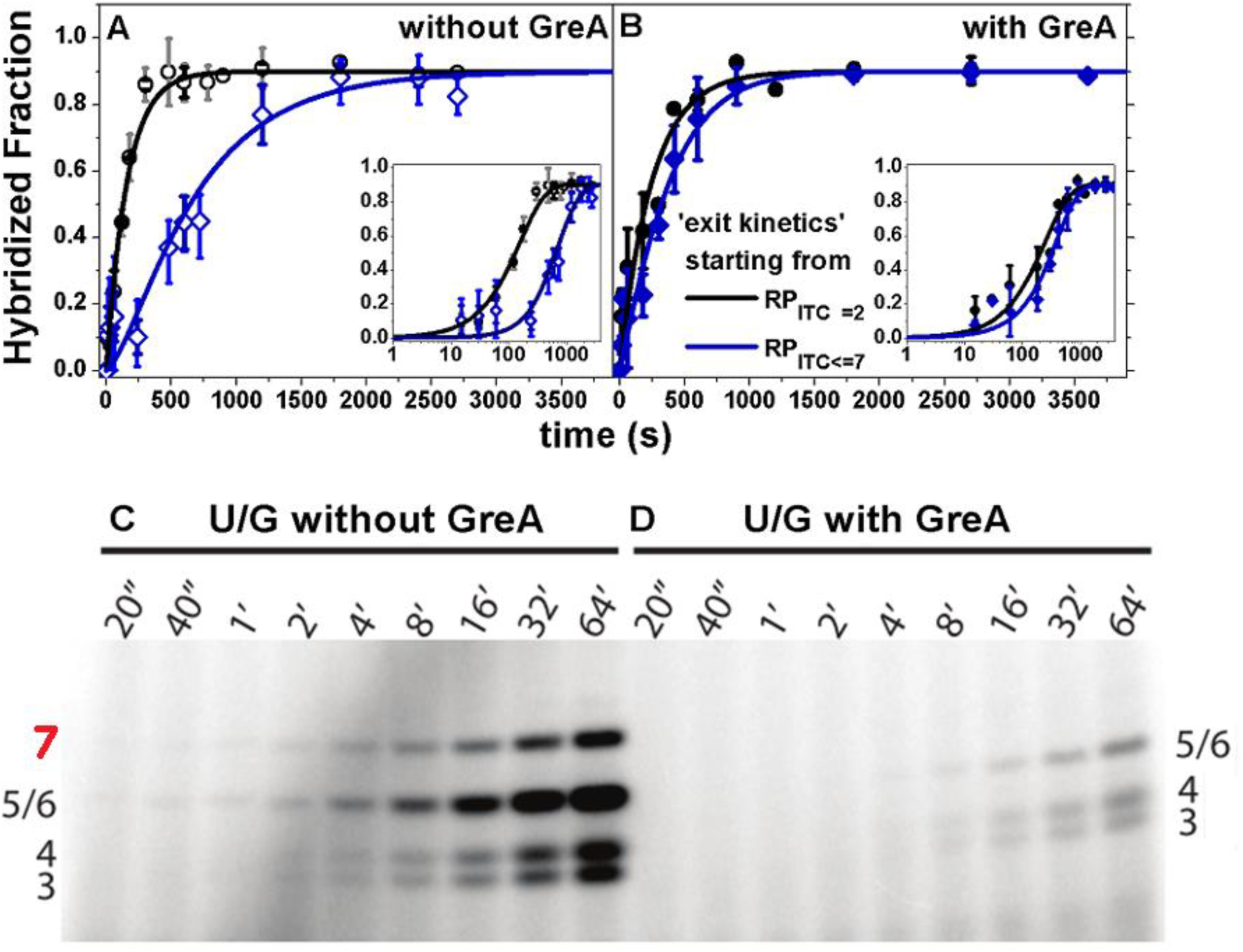
GreA rescues the initiation-related delay and cleaves backtracked RNA in initiating RNAP. (A) Run-off transcription kinetics is slower when starting from RP_ITC≤7_ (blue) than from RP_ITC=2_ (black). (B) With 1 μM GreA, the delay in transcription initiation is reduced. (C & D) Gel-based abortive initiation kinetics: without GreA, NTP-starved RP^ITC≤7^ produced abortive transcripts of up to 7 bases long, whereas this product is not produced with 1 μM GreA, suggesting 1-2 bases of 3’-backtracked RNA is cleaved by GreA during initiation.

To further test the effect of GreA during RNAP transcription initiation, we performed *in vitro* transcription assays in which ^32^P-labeled RNAP transcripts were quantitated following poly-acrylamide gel electrophoresis (PAGE). This enabled identification of various abortive initiation products (band assignment in **Fig. S4A**) and thus provided a means to determine whether GreA catalyzed cleavage of short transcripts during transcription initiation. As shown in **Figs. 2C** & **2D**, the 7-base abortive RNA product was not generated in the presence of GreA. Because GreA stimulates cleavage only of “backtracked” RNA (i.e. with the 3’-end inserted into the secondary channel), these data, combined with our single-round kinetics data, confirmed that RNAP is capable of backtracking during transcription initiation. Because 5-and 6-base transcripts co-migrate on the polyacrylamide gels, we cannot confirm whether one, two, or both one and two bases are cleaved by GreA in this assay (see Methods).

### RNAP backtracking and pausing observed with all NTPs present

Although reduced NTP levels may occur *in vivo* (e.g. metabolic stress), the absence of select NTPs is unlikely. Therefore, we next asked whether RNAP backtracking during initiation would occur under more physiologically relevant conditions. We initially examined transcription with the quenched kinetics assay under NTP concentration imbalance (UTP and GTP >> ATP and CTP at the lacCONS promoter and UTP and ATP >> CTP and GTP at the T5N25 promoter). Consistent with the results described above, we observed a delay in exit kinetics from the RP_ITC=2_ state under conditions of NTP imbalance (compared to equimolar conditions; **Fig. S5**) at each of the two different promoter templates tested (**Fig. S1**).

The single-round quenched kinetics and transcription assays with ^32^P-labeled UTP are ensemble experiments, which cannot reliably detect rare events. Indeed, with all NTPs in equimolar amounts (100 μM), we did not observe the 7-base abortive transcript, suggesting that when NTPs are abundant, RNAP rarely backtracks from the RP_ITC≤7_ state (**Fig. S4B**). To detect potentially rare intermediates under more physiologically relevant conditions of equimolar NTPs, we implemented magnetic tweezers experiments with supercoiled promoter templates (**Fig. 3A**). This assay allowed us to track individual RNAP complexes over time and simultaneously detect and identify distinct RPITC states, based upon well-established changes in DNA extension (3). In the absence of GreA, we observed short‐ and long-lived RP_ITC_ states (**Figs. 3B**, **3D**, **3F** & **3H**). The lifetimes spent in RP_ITC_ states are summarized in a histogram fitted with a double exponential in which 90% of events (n=216) were short-lived (τ = 300±40 s SEM), and 10% were long-lived (τ = 2600±700 s SEM; **Fig. 3H, blue**). Correlating these data with DNA bubble sizes (representing distinct RP_ITC_ states; **Figs. 3A** & **3F**) revealed that 90% of events were characterized by a 15 ± 3 bp transcription bubble. The remaining 10% of events were characterized by a more homogeneous 10 ± 1 bp transcription bubble. In agreement, correlative analyses (**Fig. 3D**) indicated RP_ITC_ states with a shorter bubble (<12 bases) can be long-lived (~2200 ± 350 s SEM, n=37;**Fig. 3D oval**) whereas RP_ITC_ states with a larger bubble (>12 bases) were generally shorter-lived (~1100 ± 110 s SEM, n=179).

**Fig. 3:**
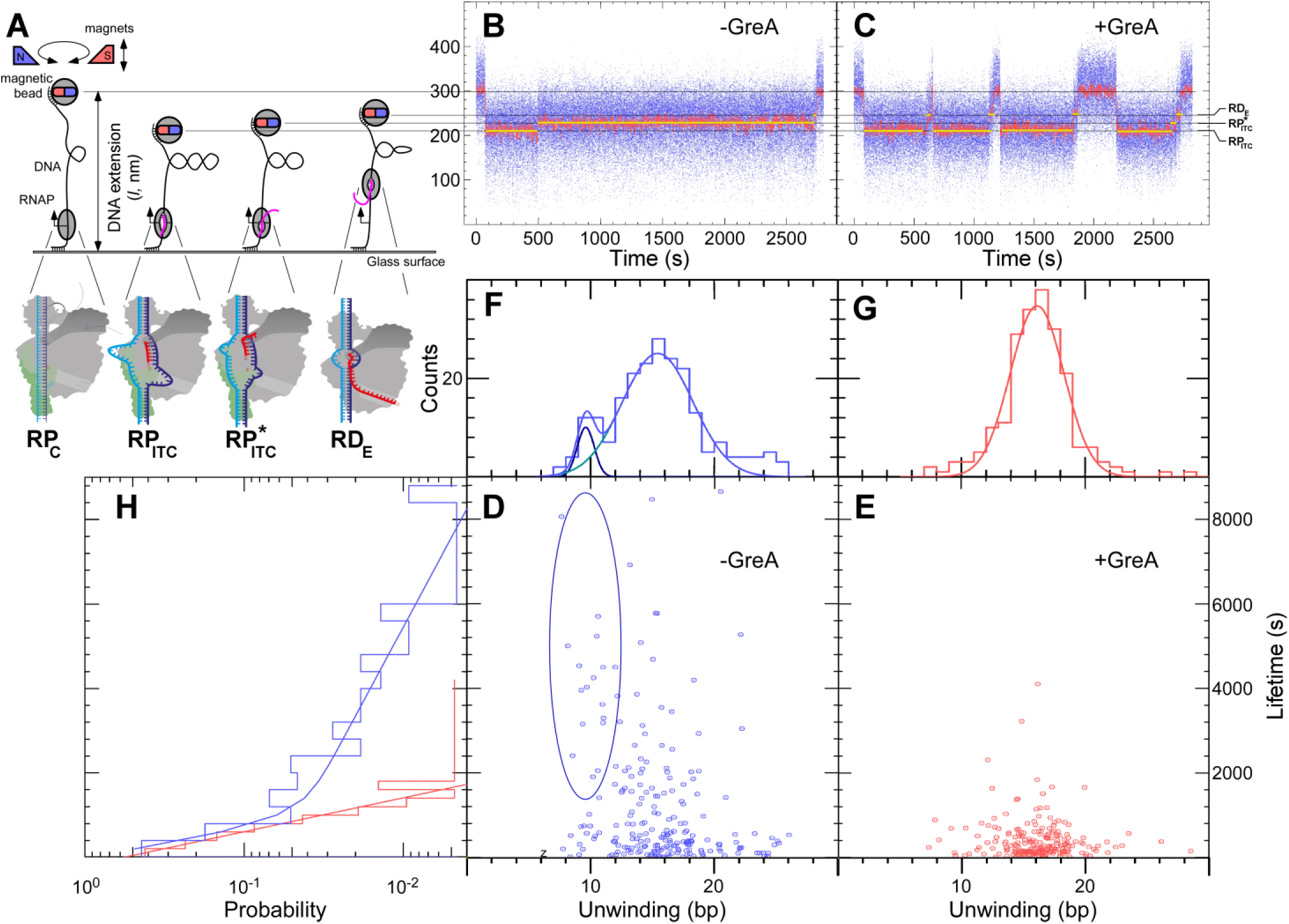
Backtracking in initiation correlates with RNAP pausing in the presence of equimolar NTPs. (A) Schematics of the magnetic tweezer transcription assay (see Materials and Methods). Representative bead extension trajectories shown for single-molecule transcription experiments without (B) or with (C) 1 μM GreA unwinding levels (grey lines) shown, indicating different bubble sizes imposed by different RNAP states (below). Yellow lines highlight typical lifetimes in each state. Unwinding levels and RP_ITC_ and RP*_ITC_ lifetimes are summarized in scatter plots without (D) or with (E) 1 μM GreA; their 1D projections are shown in (F), (G), and (H). Ellipse in (D) highlights events with long lifetimes and reduced unwinding (see also in (F)), that are absent with GreA (E, G). 20-50 DNA templates used for each condition, with 5-10 transcription pulses per template.

These data suggested that, in addition to the well-characterized RPITC state (**Fig. 3A**), a subset of RNAP complexes entered a distinct, long-lived state characterized by a smaller transcription bubble (denoted RP*_ITC_ in **Fig. 3A**, **3B** & **3C**). We hypothesized that this long-lived initiation intermediate represented backtracked RNAP previously characterized in our quenched kinetics and gel-based transcription assays. If correct, the addition of GreA would be expected to markedly reduce the number of these long-lived events. In agreement, RP_ITC_ states (n=209) became uniformly short-lived (τ = 350 ± 30 s SEM) in the presence of GreA, with transcription bubbles of larger sizes (16 ± 2 bases (SEM); **Figs. 3C**, **3E**, **3G** & **3H**). Experiments completed on a different promoter template showed the same trend (**Fig. S6**). Thus, the presence of GreA increased the number of events associated with a larger bubble size and caused a net reduction in lifetime (**Fig. S6D & S6E, quadrants**). These data further support the existence of a long-lived backtracked state during initiation and reveal a role for GreA in preventing RNAP from entering this state.

## Conclusions

Our results support the existence of a previously uncharacterized state in which RNAP backtracks and pauses during transcription initiation; moreover, GreA and NTP availability appear to play key roles in regulating the flux in or out of this state. Because basic mechanisms of transcription by RNA polymerases are broadly conserved (21) (e.g. scrunching (2, 3, 22), similar mechanistic role of σ70 and TFIIB (23, 24), trigger-loop function in catalysis (25–29)), it will be important to determine whether similar backtracked states are adopted during transcription initiation by eukaryotic RNA polymerases. Mammalian RNA polymerase II (Pol II) pauses during early stages of transcription, and this represents a common regulatory intermediate (30). Potentially, a mechanistic intermediate of paused mammalian Pol II enzymes may involve RNA backtracking; such backtracked intermediates may help explain why TFIIS, a eukaryotic ortholog of GreA, has been linked to transcription initiation and assembles with Pol II at the promoter (31–33). Finally, we emphasize that, because RNAP transcription initiation is rate-limiting and highly regulated *in vivo*, this previously unidentified backtracked/paused RNAP state may lead to potential new strategies for molecular therapeutics and to the development of novel antibiotics.

## Methods

### 1. Transcription quenched kinetics assay

#### 1.1 Preparation of a stable RP_O_, RP_ITC=2_

RP_O_ solution is prepared with 3 μL *E. coli* RNAP holoenzyme (NEB, Ipswich, MA, USA, M0551S; 1.6 μM), 10 μL 2X transcription buffer (80 mM HEPES KOH, 100 mM KCl, 20 mM MgCl_2_, 2 mM dithiotreitol (DTT), 2 mM 2-mercaptoethylamine-HCl (MEA), 200 μg/mL Bovine Serum Albumin (BSA), pH 7), 1 μL of lacCONS+20A promoter(16) (sequence in **Figs. 1A & S1**) and 6 μL of water. RP_O_ is then incubated in solution at 37^0^C for 30 minutes. To remove nonspecifically-bound RNAP, 1 μL of 100 mg/mL Heparin-Sepharose CL-6B beads (GE Healthcare, Little Chalfont, Buckinghamshire, UK) is added to RP_O_ solution together with 10 μL of pre-warmed 1X transcription buffer. The mixture is incubated for 1 minute at 37^0^C and centrifuged for at least 45 seconds at 6000 rpm. 20 μL of the supernatant containing RP_0_ formed on lacCONS or T5N25 promoters (sequences in **Figs. 1A & S1**) are transferred into a new tube for an extra incubation with 1.5 μL of 10 mM Adenylyl(3′-5′) adenosine or Adenylyl(3′-5′)uridine (A_p_A or A_p_U; Ribomed, Carlsbad, CA, USA) at 37^0^C for 20 minutes, respectively, to form RP_ITC=2_ solutions. These RP_ITC=2_ solution s are used as stock for all transcription reactions. 2 μL of RNAse inhibitor (NEB, Ipswich, MA, USA, M0314S) are added into the RP_ITC=2_ solution to prevent degradation of newly synthesized RNA molecules.

#### 1.2 Design and measurement of the transcription kinetics

To produce run-off transcripts, high-purity ribonucleotide triphosphates (NTPs) (GE Healthcare, Little Chalfont, Buckinghamshire, UK) were used in all transcription reactions at 100 μM each. To obtain a specific initiation or elongation state, only a partial set of NTPs was used. The choice of the partial set of NTPs depended on the sequence of the coding region of the nontemplate strand of the promoter used (lacCONS and T5N25, see **Figs. 1A & S1**) and on nucleotide position at position +3 relative to the transcription start site (TSS). The presence of A_p_A (in lacCONS) or A_p_U (in T5N25) in RP_ITC=2_ stabilized RNA up to position (+2), but also prevented transcription initiation from an unwanted site(34) and diversion into an unproductive pathway(35, 36)). The NTP starvation schemes depend on the different initially transcribed sequences (ITS) being used (**Figs. 1A & S1**). To exit from the initiation/elongation NTP-starved state the reaction mixture was complemented with all four NTPs.

For kinetics, the reaction mixture is incubated with the partial set of NTPs for a constant duration of 40 minutes at 37^0^C. The missing NTPs are then added to the reaction mixture and incubated for different durations (which make up the samples for the different timepoints in the kinetics) at 37^0^C, at which 0.5M Guanidine Hydrochloride (GndHCl) is added to quench the reaction. Subsequently, a ssDNA FRET probe is added and hybridizes with the target run-off transcript (see **Figs. 1A & S1** for probe target sequence). To confirm that reaction kinetics are not affected by changes in pH, we measured the pH of the solution before and after quenching and found that it did not deviate much from the pH 7 of the buffer used (6.8 – 7.0). For transcription kinetics experiments with GreA, 1 μM of protein factor is added to transcription complexes in NTP-starved initiation or elongation states.

After quenching a transcription reaction with 0.5 M GndHCl, 100 μM of ssDNA FRET probe was added and incubated with the quenched reaction mixture for an additional 20 minutes at room temperature. The quenched-probed reaction mixtures were then used for μsALEX measurements. An example of the quenched kinetic assay FRET results is shown in **Fig. 1C**.

After treatment with Heparin-Sepharose beads, the exact RNAP-promoter complex concentration is unknown, because some of the complexes had RNAP bound non-specifically to the promoter DNA. In addition, the activity of RNAP, dictated by the fraction of RNAP-promoter complexes that yield a full transcript, changes depending on various conditions. Therefore, the concentration of the DNA-RNAP complexes is calibrated beforehand to yield a dynamic range of low FRET population-fraction between 0 and 0.9 with each change in conditions. Because the concentration of promoter DNA remains unchanged after the treatment with Heparin Sepharose beads, based on the promoter DNA concentration (1 nM), we estimate that less than 1 nM of RNAP-DNA complexes are used for all *in-vitro* single-round quenched kinetics assays.

The last 20 bp of all transcribed DNA sequences code for a RNA with a stretch of 20 A (**Figs. 1C** & **S1**). The sequence of the complementary ssDNA FRET probe is 20 dT and it is doubly end-labeled with a pair of fluorophores suitable for smFRET: a donor, Tetramethylrhodamine, at the 5’-end (5’ TAMRA modification), and an acceptor, Alexa Fluor 647, at the 3’-end (3’ Alexa Fluor 647 modification); ordered from IDT, Coralville, IA, USA ((16)).

Each time point in the quenched kinetics assay is measured for a duration of 10-15 minutes using a setup described in Panzeri et. al. ((37)) using Perkin Elmer SPADs and 532 and 638 nm CW lasers operating at powers of 170 and 80 μW, respectively.

Each kinetic measurement was performed at least in duplicates using different preparations obtained on different days. For each batch, we made sure that:

1. The FRET probe in the presence of RP_ITC=2_ without NTPs yielded only a high FRET population (negative control).
2. The kinetic trace reaches a hybridized fraction (low FRET sub-population) of 0.90±0.05.
3. After a long incubation of RP_ITC=2_ with all 4 NTPs (20 minutes), the fraction of hybridized probe reaches 0.90±0.05 (positive control). This control is performed daily on the same batch used to prepare NTP-starved RNAP states.
4. After a very long incubation time (typically several hours) of a sample with a quenched reaction, the measurement yielded the same low FRET population-fraction (quenching does work)

The negative control measurement yields a single high FRET efficiency population and serves as the “t=0” time point. The positive control measurement results in a 90±% of the bursts belonging to the low FRET efficiency sub-population and serves as the asymptotic kinetic value at very long times, “t=∞”.

The difference between exit kinetics from RP_ITC=2_ and exit kinetics from an NTP-starved state, both prepared from the same batch, is solely the time at which all four NTPs were added. All other experimental conditions (concentrations, temperature, etc.) are identical per batch. Therefore, any changes in activity that may be caused solely due to the starvation of NTPs will show a change in the hybridized fraction in long timepoints of the kinetic trace. Such comparisons were routinely performed and have never shown a difference in the long timepoint baseline between the two kinetics from RP_ITC=2_ and from NTP-starved states (within 5% error). Therefore, the abovementioned positive control served as a proof that experimental conditions (e.g. NTP-starvation) did not alter the relative activity.

The results shown in **Figs. 1D, 2A, 2B & S2** are all averages of such repeated measurements (examples of these repeats are shown in **Fig. S**7). The error bars reported in these figures are the standard deviation of the repeated measurements. The values at the end of the kinetic trace of repeated measurements were very close to 0.9 in all repeats. In order to compare kinetics starting from different states, however, we had to normalize all kinetic traces so that all of them end exactly at 0.9.

All quenched kinetics data were globally fit to a simplified model that allows the run-off production kinetics to go either directly from an on-pathway initiation state (*On*) to elongation (*RunOff*) or to start at an off-pathway state (the backtracked and paused state; *Off*) and then go to elongation through slow recovery to the on-pathway initiation state as in the following schematics:

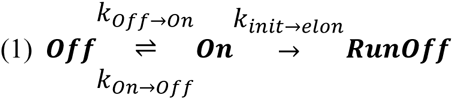

The data was globally fitted to the model assuming the on-pathway state is RP_ITC=2_, hence kinetics starting from RP_ITC=2_ is fitted with the model assuming that at t=0 all molecules are occupying the on-pathway state, while kinetics starting from RP_ITC≤4,6,7_ is fitted assuming at t=0 all molecules are occupying either the on-pathway or the off-pathway initiation states.

The results of the global fit are shown on the figures as continuous lines and the best fit values (the rate constants and the on-pathway population at t=0) are reported in **Table S1**.

#### 1.3 smFRET analysis for the quantification of transcription kinetics

Dual-color fluorescence photon-timestamps from freely diffusing molecules are recorded using an ALEX-FAMS set-up ((10, 11)). Fluorescence bursts are identified in the recorded stream of photon-timestamps, and the number of photons in a burst and the burst start/stop times are tabulated. Each burst is identified using an sliding-window burst search that looks for consecutive m(=10) photons exhibiting a count rate higher than a given threshold parameter F(=6) times the background rate ((38, 39)). The background rate is estimated as a function of time (typically over time-durations of 30s) via maximum likelihood fitting of the inter-photon delays distribution. This assures that slow changes in the background rate are accounted for. In singlemolecule μsALEX analysis, three streams of photons are analyzed: donor and acceptor fluorescence photons during green laser excitation (noted here as DD and DA, respectively), and acceptor photons during red laser excitation (noted here as AA). Burst photon counts in each of these photon streams, are background-corrected by subtracting the burst duration times the background rate. First, an all-photon (all streams) burst search is applied. After filtering for bursts with sizes larger than 25 photons, the proximity ratio and the stoichiometry are calculated for each burst to identify the sub-population of bursts where both donor and acceptor are active (FRET sub-population), sub-population of donor-only fluorescence bursts (DO), and subpopulation of acceptor-only fluorescence bursts (AO) ((40)). Next, correction factors are calculated according to Lee et al.(40). These correction factors include the donor fluorescence leakage into the acceptor detection channel (lk~0.07), factor that accounts for acceptors directly excited by the green laser (dir~0.04) and the factor correcting for differences in donor and acceptor quantum yields and detection efficiencies (γ~0.61).

Next, a dual channel burst search (DCBS; intersection of bursts from green excitation burst search and red excitation burst search) ((41)) is performed using m=10 and F=6, in order to isolate the FRET-only subpopulation for further analysis. After all correction factors are applied, the following two conditions are used to isolate smFRET data (on results of DCBS with m=10 and F=6):

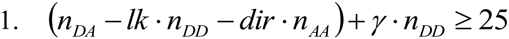

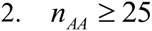

Each transcription quenched kinetics time point consists of the same two FRET populations but with different fractions that follow the evolution of the run-off transcript production (**Fig. 1C**). All corrected FRET histograms of all time points in a given kinetic trace are globally fitted to a sum of two Gaussians. In the context of the global model, the means and widths of the sub-populations are constrained to be constant as a function of time (i.e. the same for all datasets), while the amplitudes are left free to vary for each timepoint.

The background-dependent burst search and selection in this work was performed using FRETBursts, an open source burst analysis program for smFRET data (42). Model fittings were performed using Matlab scripts (MathWorks Matlab, Natick, MA, USA) through the *lsqcurvefit* nonlinear regression function.

### 2. Transcription assays visualizing abortive product formation using urea-denaturing PAGE analysis of [^32^P]-radiolabeled RNA products

Abortive transcription assays were run using the lacCONS promoter having its probe target 20A sequence replaced by the WT lacUV5 sequence at registers from +20 to +39 (**Fig. S4**). Three units of RNAP holoenzyme (NEB, Ipswich, MA, USA, M0551S) were mixed with 50 nM promoter DNA in 1x transcription buffer in a final volume of 20 μL. The reaction was then incubated at 37 ^0^C for 20 minutes to form RP_O_, followed by addition of 1 μL of 100 mg/mL Heparin-Sepharose beads and 10 μL of transcription buffer. The mixture was incubated for ~1 minute, centrifuged and 20 μL of the supernatant was removed and added to 10 μL pre-warmed transcription buffer. After incubating an additional 10 minutes, ApA was added at a final concentration of 1.3 mM and incubated for 30 minutes to form the RP_ITC=2_. The RP_ITC=2_ was then diluted to 400 μL with transcription buffer containing SUPERaseIN (AM2696, Thermo Fisher scientific, Waltham, MA, USA) to final concentrations of 1.7 nM template, 112 μM A_p_A and 0.3 units/ μL SUPERaseIN. This solution was stored at room temperature and used as a stock for each time course.

For time course experiments, 90 μL of the stock solution was briefly incubated to bring it to 37 ^0^C. To analyze the production kinetics of abortive products from RP_ITC≤7_, stock solution was mixed with 10 μL of 200 μM UTP+GTP mixture supplemented with ~ 10 μCi [α^32^P]UTP. At each time point, 10 μL aliquot was then removed and mixed with an equal volume of formamide gel loading buffer. To analyze abortive product formation from RNAP that was not stalled, the UTP+GTP mixture was replaced by a complete set of NTPs. In experiments looking at the effects of GreA on abortive product formation, an additional 15-minute incubation at 37 ^0^C was performed before the addition of NTPs, either in the presence or absence of 1 μM GreA. The stopped reaction aliquots were stored at ‐20^0^C until running the urea-denaturing PAGE.

Samples were heated for 3 minutes at 90 ^0^C and loaded on a 23%, (19:1 acrylamide:bis-acrylamide) 0.4 mm thick urea-denaturing polyacrylamide gel. The gel usually ran for 5 to 6 hours at 1500 V in 1x TBE with an additional 0.3M sodium acetate in the bottom well. The gels were then removed, dried, and exposed on a phosphor-storage screen about 2 days. Screens were visualized using a Typhoon Phosporlmager.

### 3. Magnetic trapping assay

#### 3.1 DNA constructs

We designed and had custom-synthesized (Eurofins MWG) DNA fragments flanked by KpnI sites containing a modular region for insertion of a promoter and initial transcript, followed by a transcribed region and a terminator. The modular region is flanked by HindIII and SpeI sites:

5’ GGTACC<u>AAGCTT</u>GCGAACTGCACTCGGAAC**<u>ACTAGT</u>**ATGCATCGAATAGCCATCCCAATCGATATCGAGGAGTTTAAATATGGCTGATGCATGAATTCGTTAATAACAGGCCTGCTGGTAATCGCAGGCCTTTTTATTTGGGAATTCGGTACC
where Kpnl sites are indicated in red; HindIII and SpeI sites are underlined; and the tR2 terminator is in purple. This transcription backbone was cloned into the KpnI site of the *Th. aquaticus* RPOC gene, and a 2.2 kbp subfragment of this construct centered about the transcription unit was PCR amplified and subcloned into the XbaI and SbfI sites of pUC18 using HiFi thermostable polymerase (Roche) and PCR primers (XbaI and SbfI sites underlined):

5’ GAGAGATCTAGAGACCTTCTGGATCTCGTCCACCAGGand5’ GAGAG<u>ACCTGCAGGA</u>CATCAAGGACGAGGTGTGG

We then cloned the lacCONS promoter into the HindIII and SpeI sites underlined above using the oligo-based dsDNA fragment with the top strand:

5’ AGCTAGGCTTGACACTTTATGCTTCGGCTCGTATAATGTGTGGAATTGTGAGAGCGGATTAG

Similarly, we cloned the T5N25 promoter using the dsDNA fragment with the following top strand:

5’ AGCTAAAAATTTATTTGCTTTCAGGAAAATTTTTCTGTATAATAGATTCATAAATTTGAGAGAGGAGTCC

DNA for single-molecule experiments was prepared from freshly grown DH5α by ion-exchange chromatography (Macherey-Nagel), digested with XbaI and SbfI, and the 2.2 Kb band isolated by gel purification and extraction using spin column (Macherey Nagel).

The 2.2 kbp DNA fragments containing the centrally-located transcription unit were ligated at the XbaI site to 1 kbp DNA multiply-labelled with biotin, and at the SbfI site to 1 kbp DNA multiply-labelled with digoxigenin. Labelled DNAs were synthesized via PCR carried out in the presence of dUTP-biotin and dUTP-digoxigenin, respectively (Roche) (43, 44).

#### 3.2 Single-Molecule experiments

Functionalized 2.2 kbp DNA molecules were first attached to 1 μm-diameter streptavidin-coated magnetic beads (MyOne Streptavidin C1, Life Technologies), and then tethered to a modified glass capillary surface coated with anti-digoxigenin (Roche) (44). Experiments were carried out on a homemade magnetic tweezer microscope to extend and supercoil the DNA, running the PicoTwist software suite to track and analyze the position of the magnetic bead. This position marks the free end, and thus the extension of the functionalized DNA. Data were analyzed using custom routines in the Xvin software suite.

In the supercoiling transcription assay where plectonemic supercoils are present (+4 positive supercoils throughout), the extension changes of the DNA construct report on the number of supercoils. Specifically, the DNA typically contracts by ~55 nm for every additional supercoil when extended at low force (F=0.3 pN) as in these experiments. DNA unwinding by RNAP is sensitively reported via its effect on overall DNA supercoiling: conservation of linking number means that topological unwinding of 10.5 bp results in a ~55 nm decrease in DNA extension.

Experiments were carried out in standard buffer at 34^0^C using 100 μM RNAP saturated with σ70 (prepared as in (45)) and 100 μM A_p_A (for experiments on lacCONS promoter; we used 100 μM A_p_U for experiments on T5N25 promoter) and 100 μM each of ATP, UTP, GTP and CTP. When added, GreA is at 1 μM.

### 4. Illustrations

All illustrations of RNAP transcription initiation and elongation states have been prepared in Adobe Illustrator CC 2015 (San Jose, CA, USA).

## Acknowledgements

Author Contributions: E.L. and S.C. developed and designed the quenched-kinetics experiments. E.L., S.C., W.S.L., W.L.G., and Y.A. performed the quenched-kinetics measurements, in consultation with A.I. and X.M. B.A. designed and performed gel-based experiments; S.W. and T.S. designed, performed and analyzed magnetic tweezers experiments; J.J.L. and S.B. prepared GreA protein. E.L., S.C., B.A., S.B., D.J.T., and S. W. analyzed the data; E.L., S.C., T.S., B.A., D.J.T., and S.W. wrote the paper.

We thank Prof. William Gelbart, Prof. Charles Knobler, Dr Cathy Yan Jin and Xiyu Yi for fruitful discussions and Maya Lerner for preparation of illustrations. This work was funded by the NIH (GM069709 to SW, GM095904 to XM and SW) and NSF (MCB-1244175 to SW and DJT).

## References

1. Straney DC & Crothers DM (1987) A stressed intermediate in the formation of stably initiated RNA chains at the Escherichia coli lac UV5 promoter. Journal of molecular biology 193(2):267–278.

2. Kapanidis AN, et al. (2006) Initial transcription by RNA polymerase proceeds through a DNA-scrunching mechanism. Science 314(5802): 1144–1147.

3. Revyakin A, Liu C, Ebright RH, & Strick TR (2006) Abortive initiation and productive initiation by RNA polymerase involve DNA scrunching. Science 314(5802): 1139–1143.

4. Pupov D, Kuzin I, Bass I, & Kulbachinskiy A (2014) Distinct functions of the RNA polymerase sigma subunit region 3.2 in RNA priming and promoter escape. Nucleic acids research 42(7):4494–4504.

5. Zuo Y & Steitz TA (2015) Crystal Structures of the E. coli Transcription Initiation Complexes with a Complete Bubble. Molecular cell 58(3):534–540.

6. Samanta S & Martin CT (2013) Insights into the mechanism of initial transcription in Escherichia coli RNA polymerase. The Journal of biological chemistry 288(44):31993–32003.

7. Carpousis AJ & Gralla JD (1980) Cycling of ribonucleic acid polymerase to produce oligonucleotides during initiation in vitro at the lac UV5 promoter. Biochemistry 19(14):3245–3253.

8. Hsu LM (2002) Promoter clearance and escape in prokaryotes. Biochimica et biophysica acta 1577(2): 191–207.

9. Margeat E, et al. (2006) Direct observation of abortive initiation and promoter escape within single immobilized transcription complexes. Biophysical journal 90(4): 1419–1431.

10. Kapanidis AN, et al. (2004) Fluorescence-aided molecule sorting: analysis of structure and interactions by alternating-laser excitation of single molecules. Proceedings of the National Academy of Sciences of the United States of America 101(24):8936–8941.

11. Kapanidis AN, et al. (2005) Alternating-laser excitation of single molecules. Accounts of chemical research 38(7):523–533.

12. Förster T (1948) Ann. Phys. 437:55–75.

13. Deniz AA, et al. (1999) Single-pair fluorescence resonance energy transfer on freely diffusing molecules: observation of Forster distance dependence and subpopulations. Proceedings of the National Academy of Sciences of the United States of America 96(7):3670–3675.

14. Deniz AA, et al. (2001) Ratiometric single-molecule studies of freely diffusing biomolecules. Annual review of physical chemistry 52:233–253.

15. Mukhopadhyay J, et al. (2001) Translocation of sigma(70) with RNA polymerase during transcription: fluorescence resonance energy transfer assay for movement relative to DNA. Cell 106(4):453–463.

16. Kim S, et al. (2011) High-throughput single-molecule optofluidic analysis. Nature methods 8(3):242–245.

17. Borukhov S, Lee J, & Laptenko O (2005) Bacterial transcription elongation factors: new insights into molecular mechanism of action. Molecular microbiology 55(5): 1315–1324.

18. Toulme F, et al. (2000) GreA and GreB proteins revive backtracked RNA polymerase in vivo by promoting transcript trimming. The EMBO journal 19(24):6853–6859.

19. Laptenko O, Lee J, Lomakin I, & Borukhov S (2003) Transcript cleavage factors GreA and GreB act as transient catalytic components of RNA polymerase. The EMBO journal 22(23):6322–6334.

20. Stepanova E, et al. (2007) Analysis of promoter targets for Escherichia coli transcription elongation factor GreA in vivo and in vitro. Journal of bacteriology 189(24):8772–8785.

21. Werner F & Grohmann D (2011) Evolution of multisubunit RNA polymerases in the three domains of life. Nature reviews. Microbiology 9(2):85–98.

22. Fazal FM, Meng CA, Murakami K, Kornberg RD, & Block SM (2015) Real-time observation of the initiation of RNA polymerase II transcription. Nature 525(7568):274–277.

23. Murakami KS & Darst SA (2003) Bacterial RNA polymerases: the wholo story. Current opinion in structural biology 13(1):31–39.

24. Sainsbury S, Niesser J, & Cramer P (2013) Structure and function of the initially transcribing RNA polymerase II-TFIIB complex. Nature 493(7432):437–440.

25. Allison LA, Moyle M, Shales M, & Ingles CJ (1985) Extensive homology among the largest subunits of eukaryotic and prokaryotic RNA polymerases. Cell 42(2):599–610.

26. Bar-Nahum G, et al. (2005) A ratchet mechanism of transcription elongation and its control. Cell 120(2): 183–193.

27. Hekmatpanah DS & Young RA (1991) Mutations in a conserved region of RNA polymerase II influence the accuracy of mRNA start site selection. Molecular and cellular biology 11(11):5781–5791.

28. Thuillier V, Brun I, Sentenac A, & Werner M (1996) Mutations in the alpha-amanitin conserved domain of the largest subunit of yeast RNA polymerase III affect pausing, RNA cleavage and transcriptional transitions. The EMBO journal 15(3):618–629.

29. Weilbaecher R, Hebron C, Feng G, & Landick R (1994) Termination-altering amino acid substitutions in the beta' subunit of Escherichia coli RNA polymerase identify regions involved in RNA chain elongation. Genes & development 8(23):2913–2927.

30. Adelman K & Lis JT (2012) Promoter-proximal pausing of RNA polymerase II: emerging roles in metazoans. Nature reviews. Genetics 13(10):720–731.

31. Guglielmi B, Soutourina J, Esnault C, & Werner M (2007) TFIIS elongation factor and Mediator act in conjunction during transcription initiation in vivo. Proceedings of the National Academy of Sciences of the United States of America 104(41):16062–16067.

32. Kim B, et al. (2007) The transcription elongation factor TFIIS is a component of RNA polymerase II preinitiation complexes. Proceedings of the National Academy of Sciences of the United States of America 104(41): 16068–16073.

33. Stepanova EV, Shevelev AB, Borukhov SI, & Severinov KV (2009) [Mechanisms of action of RNA polymerase-binding transcription factors that do not bind to DNA]. Biofizika 54(5):773–790.

34. Robb NC, et al. (2013) The transcription bubble of the RNA polymerase-promoter open complex exhibits conformational heterogeneity and millisecond-scale dynamics: implications for transcription start-site selection. Journal of molecular biology 425(5):875–885.

35. Kubori T & Shimamoto N (1996) A branched pathway in the early stage of transcription by Escherichia coli RNA polymerase. Journal of molecular biology 256(3):449–457.

36. Susa M, Sen R, & Shimamoto N (2002) Generality of the branched pathway in transcription initiation by Escherichia coli RNA polymerase. The Journal of biological chemistry 277(18): 15407–15412.

37. Panzeri F, et al. (2013) Single-molecule FRET experiments with a red-enhanced custom technology SPAD. Proceedings of SPIE‐‐the International Society for Optical Engineering 8590.

38. Michalet X, et al. (2013) Development of new photon-counting detectors for single-molecule fluorescence microscopy. Philos TR Soc B 368(1611).

39. Eggeling C, et al. (2001) Data registration and selective single-molecule analysis using multi-parameter fluorescence detection. Journal of biotechnology 86(3): 163–180.

40. Lee NK, et al. (2005) Accurate FRET measurements within single diffusing biomolecules using alternating-laser excitation. Biophysical journal 88(4):2939–2953.

41. Nir E, et al. (2006) Shot-noise limited single-molecule FRET histograms: comparison between theory and experiments. The journal of physical chemistry. B 110(44):22103–22124.

42. Ingargiola A, Lerner E, Chung S, Weiss S, & Michalet X (2016) FRETBursts: Open Source Burst Analysis Toolkit for Confocal Single-Molecule FRET. bioRxiv.

43. Revyakin A, Ebright RH, & Strick TR (2004) Promoter unwinding and promoter clearance by RNA polymerase: detection by single-molecule DNA nanomanipulation. Proceedings of the National Academy of Sciences of the United States of America 101(14):4776–4780.

44. Revyakin A, Ebright RH, & Strick TR (2005) Single-molecule DNA nanomanipulation: improved resolution through use of shorter DNA fragments. Nature methods 2(2): 127–138.

45. Howan K, et al. (2012) Initiation of transcription-coupled repair characterized at single-molecule resolution. Nature 490(7420):431–434.

